# Molecular evolution and structural analyses of spike protein COVID-19 variants in Negeri Sembilan of peninsular Malaysia

**DOI:** 10.1101/2022.12.15.520679

**Authors:** Shuhaila Mat-Sharani, Danish A/L Kumareahsan, Ismatul Nurul Asyikin Ismail, Muhamad Arif Mohamad Jamali, Liyana Azmi

## Abstract

The sharing of COVID-19 sequences worldwide has allowed for comprehensive and real-time analyses of COVID-19 genomic diversity at regional levels. Temporal distribution of COVID-19 variants and lineages enables better infection control, surveillance, and facilitates policy making for public health. 417 sequences extracted from all COVID-19 cases in Negeri Sembilan of peninsular Malaysia from July 2021 until May 2022 were used for this study. Phylogenomics revealed a total of 20 circulating lineages, of which seven are still circulating. The majority (60.4%) of viruses in Negeri Sembilan are of GRA lineage with strong representation from the Malaysian lineage BA.1.1 (24.7%). A time series analysis showed a change in the dominating circulating lineage from AY.79 to BA.1.1, which correlated to the relaxing of lockdowns implemented by the Malaysian government. Several Malaysian sub-lineages (BA.2.40.1, BA.2.57 and BA.2.9) have emerged from April 2022 onwards. Evolutionary mutations of the sub-lineages also gave rise to novel single nucleotide polymorphisms (SNPs) in the spike proteins. Out of the 70 SNPs isolated from all samples, *in silico* prediction revealed five novel SNPs that could cause functional defects to the spike protein, which are S221L, L226S, V826L, C1240F and C1243F. Structural modelling of the V826L showed that the L826 possibly confers an increase in protein flexibility within the S2 region of S protein, which supports our *in-silico* predictions.

## Introduction

In December 2019, a novel coronavirus infected a cluster of patients in Wuhan, China. With the airborne transmission of the virus SARS-CoV-2, infections spread rapidly and were facilitated by frequent travelling across continents. By January 2020, the World Health Organization (WHO) declared a public health emergency of international concern (1). From January 2020 until 5th December 2022, 641 million cases worldwide were logged, with 6 million deaths reported (WHO, 2022). Malaysia, a southeast Asian country, was not excepted during the pandemic. The first COVID-19 case in Malaysia was reported on 25th January 2020, followed by 21 cases on consecutive days. This wave was subsequently marked as the first of the country’s total four waves of infection (3). Subsequent outbreaks were then initiated by mass gatherings and were further strained by burdened healthcare systems (4). Fortunately, with international collaborations and efforts, vaccinations have been administered worldwide. As of 29th November 2022, up to 13 million doses were administered (WHO, 2022). In Malaysia, approximately 72,000 doses have been administered since February 2021, significantly increasing the burden in healthcare settings (KKMNOW, 2022).

Infection controls in Malaysia implemented include awareness and training, infection, prevention and control practices, personal protective equipment usage, vaccination, surveillance, and management of those contracting the disease (6). Malaysia also implemented movement control orders to contain high infections. Lockdowns and movement restrictions between districts and states have proven effective in containing and controlling the spread of COVID-19 cases (7). On top of regional restrictions, Malaysia has initiated the SARS-CoV-2 genomic surveillance project to enable the identification of SARS-CoV-2 variants in Malaysia (Hazim M, 2022). The selected patients fitting the sampling criteria by the Ministry of Health would then be sequenced and deposited in the GISAID (Global Initiative on Sharing Avian Influenza Data) to enable real-time analysis and surveillance of circulating variants.

The tracking of circulating variants has been made available using several nomenclatures, including the WHO label (WHO, 2022), GISAID (10) and Phylogenetic Assignment of Named Global Outbreak LINeages (Pango lineages) (11). According to WHO, variants containing mutations that increase the infection’s transmissibility and severity and escape the immune response or diagnostics are classified as variants of concern (VOC). Variants which contain mutations but are only predicted to increase the transmissibility severity of infection escapes the immune response or diagnostics are classified as variants of interest (VOI). Most of the circulating variants accounting for global infections include VOCs of Alpha, Beta, Gamma, Delta, and Omicron variants. The GISAID lineage systems are based on shared marker mutations, which comprise 11 clades. The base clade G splits up to GH/GV/GK/GR/GRY/GRA clades – all of which share the common D614G mutation known to increase viral fitness (12). The Pango nomenclature integrates genetic and geographical information, which consists of an alphabetical and numerical suffix. For example, the Alpha Beta Delta and Omicron corresponded to Pango lineages B.1.1.7, B.1.351, B.1.617.2, and B.1.1.529 (including descendent lineages). The Pango nomenclature allows the user to derive detailed outbreak information and track the evolution of lineages between regions.

Most mutations causing the evolution of SARS-CoV-2 transmissibility and virulence are clustered in the spike (S) protein. Several factors contribute to the high mutation rates, including the highly selective pressures from interacting with the host angiotensin converting enzyme2 (ACE2) and antibodies (11). The S protein consists of two subunits: S1 and S2 (13). S1 primarily involves target recognition and binding, while S2 initiates membrane fusion and endosomal escape (14). Studies have shown that positive selections within S1 arise from the mutations that would contribute to the increased fitness of SARS-CoV-2 (15). Naturally, due to the high rates of interactions occurring within the S1 region, S1 is less conserved compared to S2 thus, allows for higher rates of mutations. Continuous and frequent surveillance is thus needed to monitor for novel mutations that could increase viral transmissibility and fitness and escape the immune response.

With the increased sequencing capabilities initiated in Malaysia, there is a need to analyse and assess the genetic evolution of SARS-CoV-2 at a regional level. Negeri Sembilan is next to Selangor, which lodged the highest cases of COVID-19 throughout the pandemic. The sub-urban landscape, combined with some rural areas of Negeri Sembilan, would impose different demographics of COVID-19 infections than the country. In addition, to the best of our knowledge, no studies have assessed the effect of mutations within the S proteins in Malaysia. Our work thus highlights the genetic evolution of SARS-CoV-2 in Negeri Sembilan and unravels novel mutations within the S proteins, which could give rise to more virulent SARS-CoV-2 variants.

## Methodology

### Sampling

438 SARS-CoV-2 sequences have been extracted and analysed over the past year (July 2021 - May 2022). Eight sequences were omitted since the nucleotide sequences were < 29,000 nt. A final total of 417 sequences was extracted for analysis. The reference genome of SARS-CoV-2 used in this study is hCoV-19/Wuhan/WIV04/2019 (WIV04). Gene annotation of 417 sequences was the first step of the analysis, which was done using Viral Genome ORF Reader (VIGOR) version 4.1.2 and SNPs were extracted from the GISAID database. From the GISAID database, other information such as patient information including age, sex, and originating lab for where the sample was accrued was available. The sequenced samples also reported the respective GISAID clades, and Pangolin lineages.

### Phylogenomic analysis

Multiple Sequence Alignment (MSA) is a multiple alignment program for amino acid, or nucleotide sequences was performed using MAFFT with FFT-NS-2 methods. Once the alignment was completed, the phylogenetic tree was constructed. In this work, MEGA X software is used to construct the phylogenetic tree. The phylogenetic tree was built using the UPGMA method with a Maximum Composite Likelihood (MCL) model, uniform site rates and a 1000-time bootstrap value. After MEGA X, the Interactive Tree of Life (iTOL) was used to display and manage the phylogenetic tree for more accessible and interactive visualisation.

### Frequency of mutations and selection for PredictSNP

The single nucleotide polymorphisms for all samples were extracted from the sequences and compared to the GISAID website. The *in-silico* prediction program PredictSNP was used to infer the biological effects of all the samples (16). PredictSNP is a consensus program which comprises the predictions from APP, PhD-SNP, Polyphen-1, Polyphen-2, SIFT, SNAP, nsSNPAnalyzer, and PANTHER to predict the biological effect of a single nucleotide variation. PredictSNP combines the six best performance tools previously mentioned to improve prediction. The prediction outputs include predictions from the top six prediction tools and outcomes of the mutation residue as “Neutral” or “Deleterious”. It also provides the percentage indicating the expected accuracy of the prediction.

### Structural analysis

Samples which showed deleterious effects by PredictSNP were selected for structural analysis. The extracted spike sequence was submitted for *in silico* structure modelling by SWISS-MODEL (17). All models with a maximal identity of more than 99% were selected. Structural analyses and visualisation of the models were performed on PyMOL.

## Results

### Demographic and clade assignment of samples

Negeri Sembilan has a population density of 1.13 million and has been slowly increasing since 2000 (DOSM, 2021). The most populated district is Seremban (∼380,000 citizens), followed by Jempol (∼125,000 citizens), Port Dickson (∼106,000 citizens), Tampin (∼77,000 citizens), Kuala Pilah (∼63,000 citizens), Jelebu (∼37,000 citizens) and Rembau (∼36,000 citizens). The sampling contributions from Negeri Sembilan, however, did not reflect the population densities of the districts and were as follows: Seremban (47%, *n* = 195), Tampin (35%, *n* = 146), Kuala Pilah (7%, *n* = 31), Jempol (6%, *n* = 24), Jelebu (2%, *n =* 7), Rembau (2%, n = 7), Port Dickson (1%, *n* = 6) (Fig 1A). Several health facilities contributed to the same districts to accommodate the high population density. From July 2021 to May 2022, our study showed the highest number of sequenced samples from Seremban, specifically accrued from Hospital Tunku Jaafar (35%, *n* = 146) and Health District Office (HDO) Seremban (11%, *n =* 49). Interestingly, Tampin, the fourth highest population in Seremban, contributed to the second highest sampling of this study and only accrued samples from HDO Tampin.

**Fig 1.**
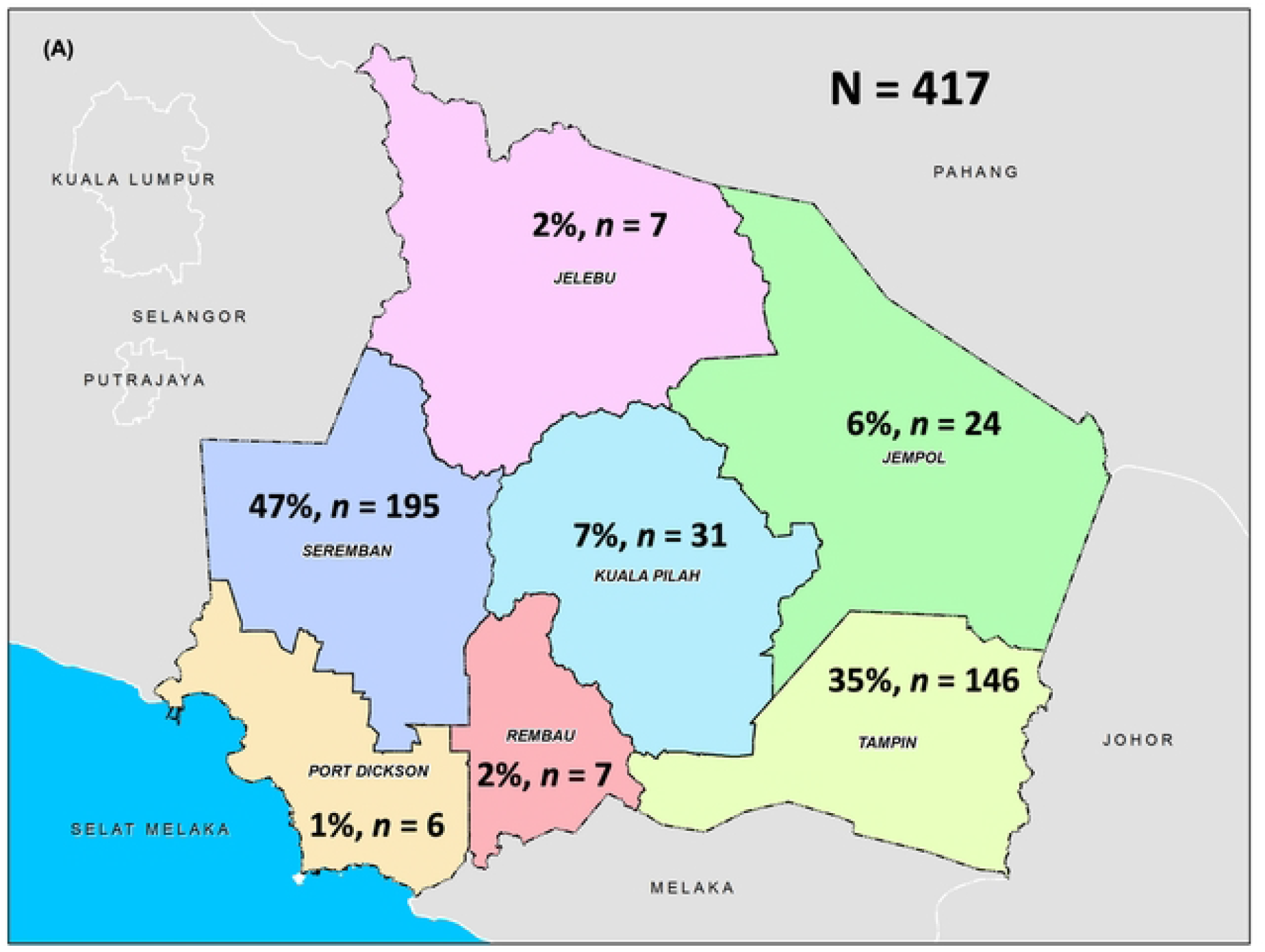
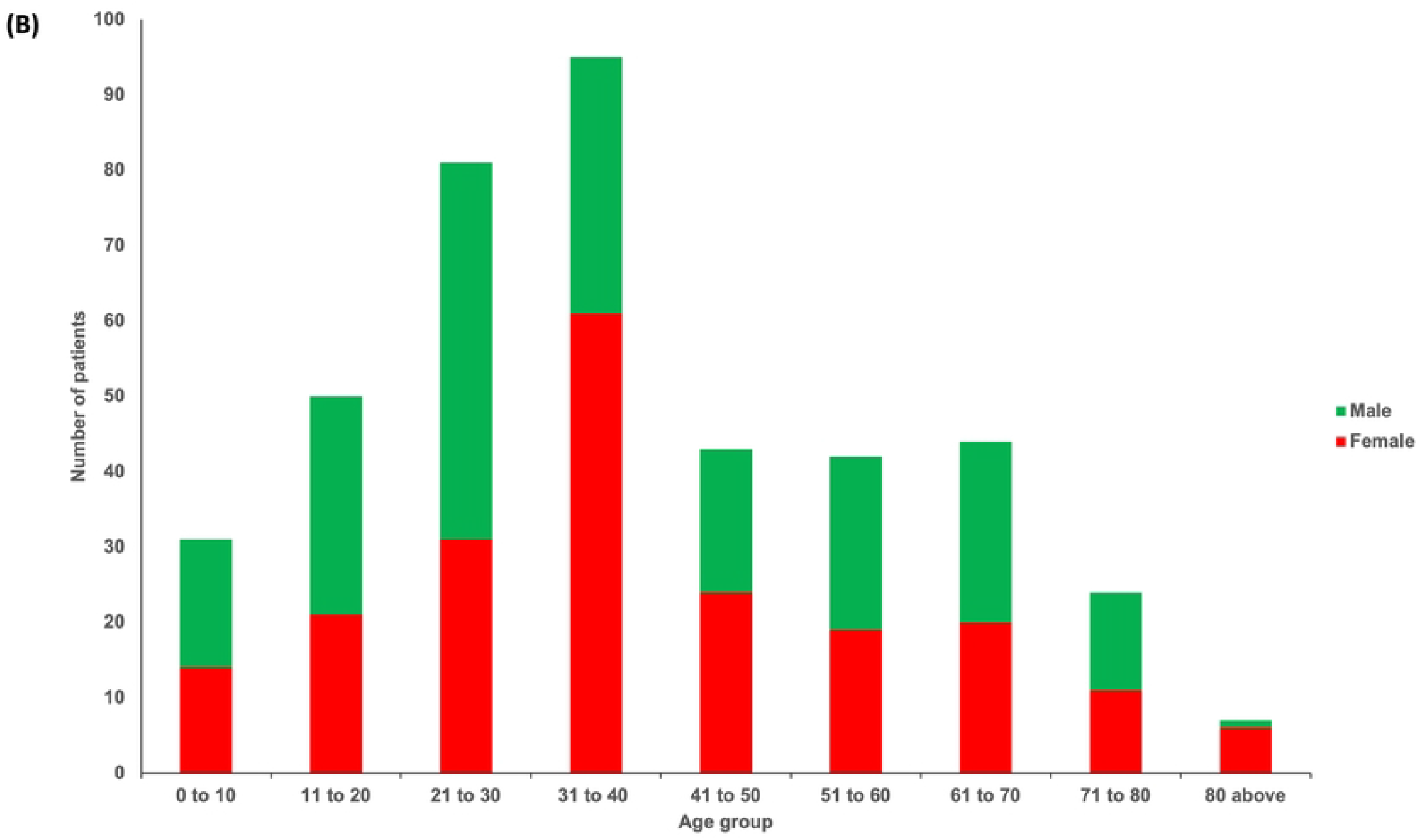
Demographic distribution of samples. (A) Regional and (B) Age distribution of samples submitted for sequencing and deposition in GISAID database.

The ratio of males to females contributing to the sampling was a 1:1, with 207 females (48%) and 210 (52%) male samples. The scattering of age across the samples was also approximately evenly distributed and as follows; 0 to10 years (33%, *n =* 33), 11 to 20 years (12%, *n =* 50), 21 to 30 years (19%, n = 81), 31 to 40 years (23%, n = 95), 41 to 50 years (10%, n = 43), 51 to 60 years (10%, n = 42), 61 to 70 years (11%, n = 44) and 80 above (1%, n = 5). Since the sampling method was universal, all data was collected and reflected the demography of COVID-19 patients from Negeri Sembilan. However, the samples also reflect the sequencing criteria set by health facilities. Therefore, criteria for sample selection for sequencing is important as specific age groups, sex, and health status, i.e., pregnancies and comorbidities, may have an impact on the virus harboured. Our study observed a 1:1 ratio of male-to-female (207:210) in the sequenced samples (Fig 1B). Age wise, the males dominated females for the age group 21-30, whereas females overwhelmed the males in the age group 31-40. Otherwise, the number of patients for both sexes are closely similar.

### Genomic epidemiology

The clade/lineage of SARS-CoV-2 was assigned based on the GISAID clade and Pango lineage assignments. From July 2021 until June 2021, the top circulating variant is Delta (61%), followed by Omicron (39%). Data showed that Delta subvariants have dominated from July 2021 to January 2022. AY.79 began circulating in Negeri Sembilan in July 2021 and persisted as the dominant variant from August 2021 until January 2022. In December 2021, BA.1.1 began circulating as the more dominant variant, but cases dropped significantly from Apr22 onwards. From January 2022, BA.2 infections began to increase and are possibly the circulating variant in Negeri Sembilan from May22 onwards. Cumulatively, the top lineage sequenced includes Delta subvariants which are AY.79 (31%, *n =* 128) and AY.59 (23%, *n =* 95). Omicron subvariants include BA.1.1 (25%, *n =* 103) and BA.2 (8%, *n =* 34) (Fig 2).

**Fig 2.**
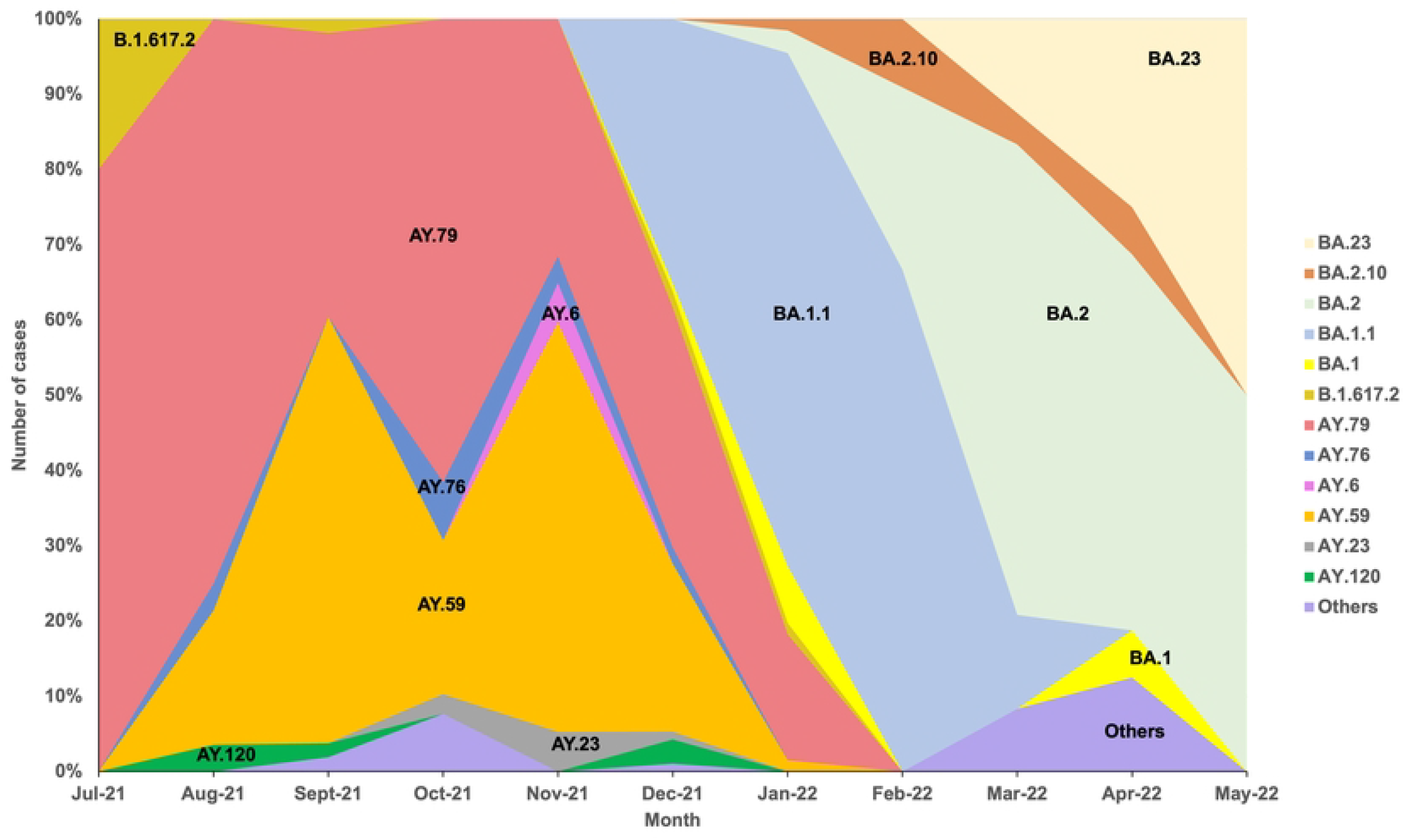
SARS-CoV-2 variants in Negeri Sembilan from July 2021 – May 2022 (n=417). Circulating SARS-CoV-2 variants in Negeri Sembilan by month. The ‘Others’ variants include AY.102, AY.42, AY.45, AY.9, BA.2.40.1, BA.2.57 and BA.2.9

### Genomic diversity of SARS-cov-2 infections in Negeri Sembilan

Of all the lineages sequenced, the Omicron variant (BA.1.1 and BA.1.1) showed the highest genetic diversity (Fig 3). The relationships of the circulating lineages have evolved, crossing all the districts in Negeri Sembilan. The phylogenetic tree of 417 revealed a total of 20 lineages from Negeri Sembilan, of which 9 (BA.1, BA.1.1, BA.1.1.18, BA.2, BA.2.10, BA.23, BA.2.40.1, BA.2.57 and BA.2.9) of the lineages are still circulating. Three major GISAID clades were identified, namely GRA, GK and O, on a maximum likelihood phylogenetic tree (Fig 3). The two major lineages from the phylogenetics tree show that mostly Delta (B.1.617.2) and Omicron (BA.1.1, BA.2.40.1, BA.2.57 and BA.2.9) variants cases highly found in Seremban and Tampin in Negeri Sembilan. The evolution of variants can clearly be seen from the O clade, followed by the GK clade, which represents Delta variants. Then, finally, the GRA clade which represents Omicron variants.

**Fig 3.**
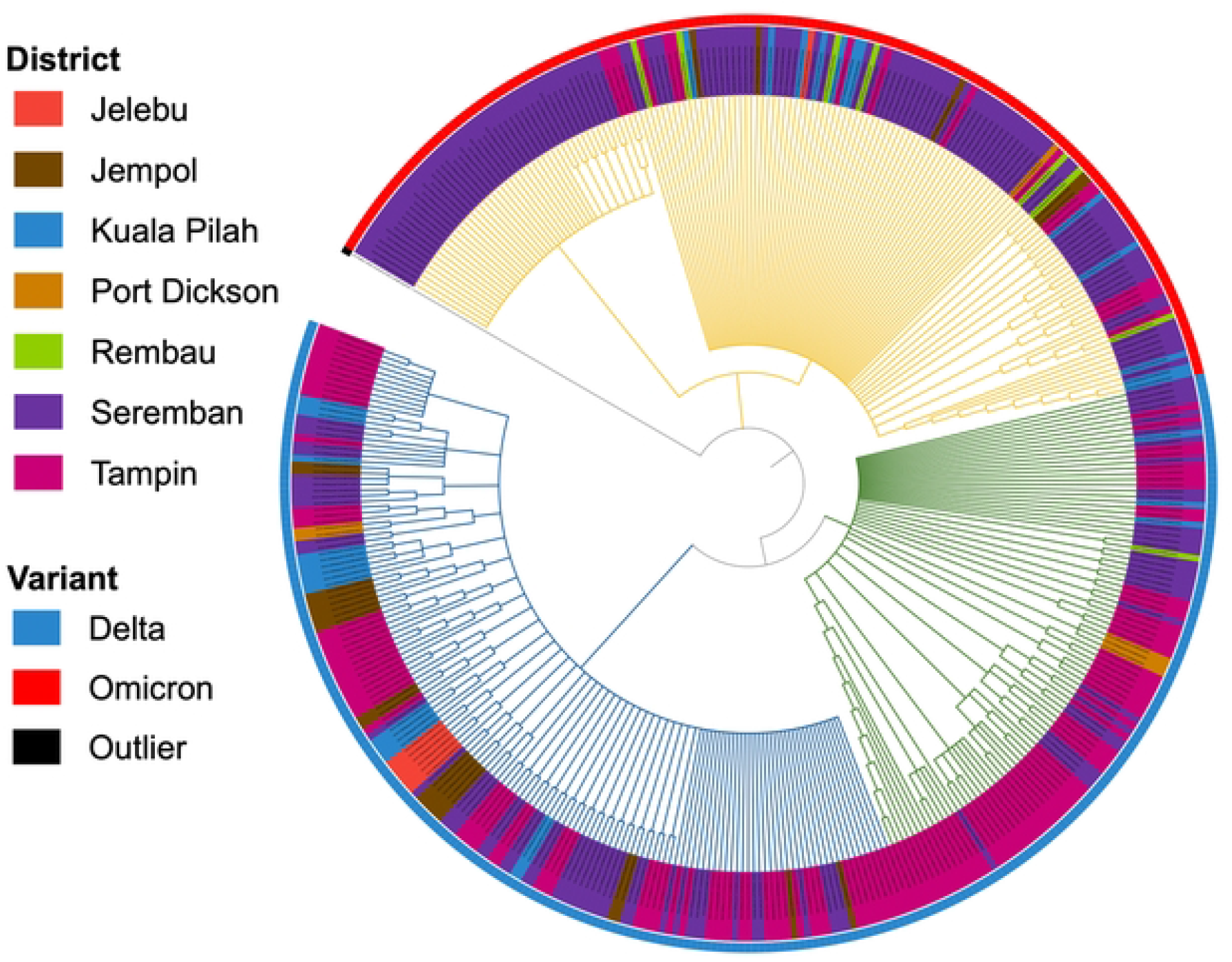
A maximum likelihood phylogenetic tree of sequenced samples in Negeri Sembilan from July 2021 – May 2022. The districts and the GISAID clades GRA (yellow), GK (green) and O (others) are as indicated.

### Frequencies of S protein SNPs

There are two dominant variants throughout sampling – AY.79 and BA.1.1. The switch between AY.79 and BA.1.1 is markedly swift, and the basis for this change was postulated to stem from several factors, including the release of lockdowns and mutations from the circulating variant. The S protein of SARS-CoV-2 is particularly a hotspot for viral mutations. It is of clinical importance due to the interaction of the S protein with immunoglobulins and the ACE-2 host cell. Mutations within the S protein have also been shown to induce immune evasion and reduce vaccine effectiveness in the community (18).

All the samples (100%) contained the D614G mutation. This mutation is included in the high frequency of isolated mutation. Other mutations with high frequency present in more than 50% of the samples, which are Y505H (57%), E156G (57%), F157del (57%), R158del (57%), T19R (59%), D950N (61%), L452R (61%), G142D (73%) and T478K (98%) mutations (Table 1). Our dataset also showed subvariants carrying rare mutations, including C1240F, D1084A, D574Y, E661D, G1251R, I1210V, I693V, L216del, L226S, N925S, P26L, Q613H, R214L, S256L, S704L, T1117I, V62I, V826L (all at 0.24%), V915I, A93S, C1243F, E1262D, L18F, N81D, V1176F, V171I, S221L (all at 0.48%), G446V, Q787K, and T573I at 1.2%, respectively.

**Table 1.** Isolated mutations with high and low frequency.

| Mutations | High frequency (N, %) |
| --- | --- |
| D614G | 414/417, 99.5 |
| T478K | 401/417, 96.4 |
| G142D | 295/417, 71 |
| L452R, D950N | 249/417, 59.9 |
| T19R | 243/417, 58.4 |
| E156G | 232/417, 55.7 |
| N501Y, P681H, H655Y, E484A | 162/417, 39 |
|  | **Low frequency (N, %)** |
| C1240F, D1084A, D574Y, E661D, G1251R, I1210V, I693V, L216del, S221L, L226S, N925S, P26L, Q613H, R214L, S256L, S704L, T1117I, V62I, V826L | 1/417, 0.24 |
| V915I, A93S, C1243F, E1262D, L18F, N81D, V1176F, V171I, S221L | 2/417, 0.48 |
| G446V, Q787K, and T573I | 5/417, 1.2 |

To understand the biological effects of these mutations, the SNPs were subjected to phenotypical predictions using PredictSNP (16). PredictSNP combines eight established predictive tools to predict the biological effect of a single nucleotide variant. The output for PredictSNP is confidence scores, with 1 being the highest score. Each submission will report as neutral (no biological change) or deleterious.

Based on the computational analysis by PredictSNP, 11 out of 417 mutations were predicted to be deleterious (S1 Table). Six mutations (T95I, N856K, N764K, N969K, Y505H, and L452R) have previously been characterised and were noted to be natural SNPs in variants of Omicron, Iota, and Mu. The other mutations were uncharacterised and showed the lowest frequency with deleterious effects, including C1240F, V826L, L226S, C1243F and S221L. C1240F, V826L and L226S. The SNPs mutations only occurred once, while C1243F and S221L mutations occurred twice (Table 2). To test the effects of these mutations on the S protein, we modelled the S protein using SWISS-MODEL with each of the sequences extracted from the respective samples.

**Table 2.** Singular SNP isolated with patient characteristics.

| Mutation | Patient characteristics |  |  |  |  |
| --- | --- | --- | --- | --- | --- |
|  | GISAID ID | Age | Gender | Lineage | Date Collected |
| C1240F | EPI_ISL_8380529 | 60 | Female | AY.59 | 13/12/2021 |
| V826L | EPI_ISL_5742653 | 51 | Female | AY.79 | 08/10/2021 |
| L226S | EPI_ISL_7649067 | 53 | Male | AY.59 | 29/11/2021 |
| C1243F | EPI_ISL_5742660 | 38 | Male | AY.4.2 | 07/10/2021 |
|  | EPI_ISL_5742690 | 38 | Female | AY.4.2 | 07/10/2021 |
| S221L | EPI_ISL_7987105 | 2 | Male | AY.79 | 22/11/2021 |
|  | EPI_ISL_8314880 | 33 | Female | AY.79 | 03/12/2021 |

### Modelling and validation of the mutant S protein

Modelling of the S mutants was done by comparing SWISS-MODEL generated structures against the reference model used for the SWISS-MODEL generated structures. The reference model is the pre-fusion SARS-CoV-2 Delta variant spike protein containing one ‘up’ RBD (ID: 7sbo.1). All selected models were made based on the highest percentage identity score to the reference model. We were interested in the SNPs located at the RBD since studies show that they could lead to better receptor engagement/immune evasion. However, all of the predicted deleterious mutations in this study were not located at the RBD. Of the five selected predicted deleterious SNPs, only V826L, S221L and L226S mutations could be visualised and modelled against the reference model (Fig 4). The C1243F and C1240F models could not be visualised due to the missing residues in the reference model. To compare the structural differences in the mutated models, the generated SWISS-MODEL was superposed onto the reference model 7sbo.1.

**Fig 4.**
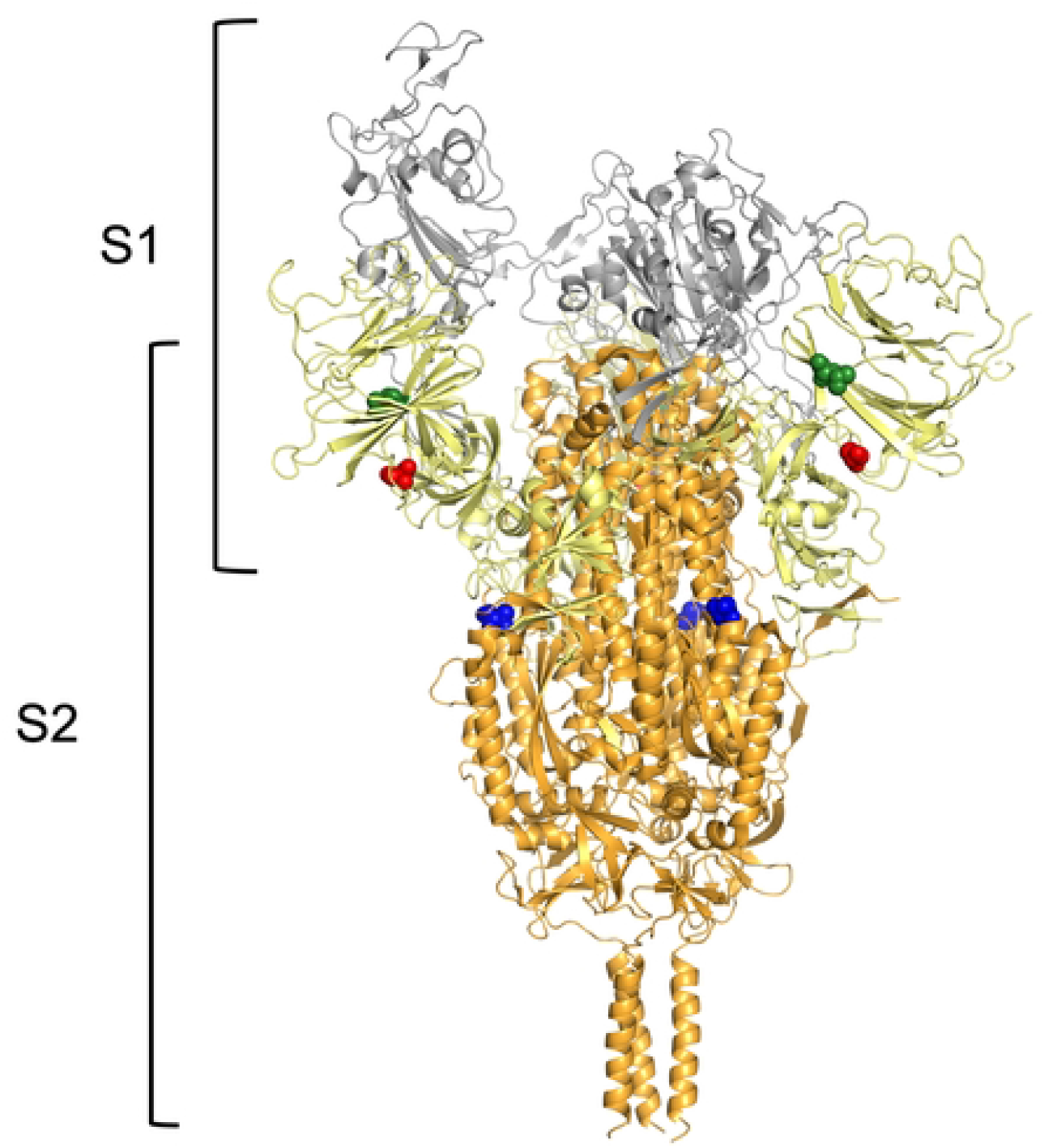
Deleterious predicted SNPs visualized on the pre-fusion SARS-CoV-2 Delta variant spike protein containing one ‘up’ RBD (ID: 7sbo.1). In the prefusion state, one of the S monomers has RBD in the ‘up’ conformation and is accessible to the receptor ACE2. The deleterious SNPs are shown in space-filled representation and reveals that V826L (blue), S221L (red) and L226 (green) are not located at the RBD site (grey).

The comparison of model S221L and L226S showed no contrasting structural changes against the reference model. However, in the V826L model, direct comparison of both structures was not available. This was due to the missing loop preceding the V826 in the reference model – presumably due to the flexibility of the loop which was visualised in the L826 model. Nevertheless, the missing loop for V826 only occurred for the S monomer in the ‘up’ conformation. Direct comparisons for V826L for the ‘down’ monomers were similar to the S221L and L226S models, and showed not significant structural differences.

## Discussion

Since the sequencing consortium initiated by the Ministry of Science, Technology, and Innovation of Malaysia, it was possible to acquire access to a higher number of sequenced genomes of COVID-19 patients in Malaysia. Work by Azami et al., (2022) (19) showed at national level, the trends for circulating SARS-COV-2 variants in the whole of Malaysia. We sought to understand if the trends hold true at regional and state level. The prevalence comparison of AY.79 of Negeri Sembilan and Malaysia shows that in Malaysia, infections by AY.79 began increasing in July 2021, reduced slightly in October 2021, and peaked in December 2021, before waning in February 2022. The highest reports of AY.79 infection were made by Terengganu and Perlis, followed by Negeri Sembilan, which suggests a route of infection that could have stemmed from Terengganu and Perlis. In addition, since the relaxing of lockdowns in October 2021, interstate travel has been available, thus allowing infection to spread between states (19). Overall, our observations were approximately similar to the trends at national level, but with a higher dominance for the AY.79 lineage. Seremban is the neighbouring district to Putrajaya and Selangor, a heavily populated district. Additionally, there was a high number of residents who reside across these districts, which will promote more movements across states and districts. The sharp spike in cases in December 2021 would correlate to the high volume of travel post-lockdowns, school holidays and mass gatherings. It was also interesting to note that since the subsiding of lockdowns in October 2021, the number of cases submitted has increased. The two significant lineages from the phylogenetics tree show that the predominating variants are Delta (B.1.617.2) and Omicron (BA.1.1, BA.2.40.1, BA.2.57 and BA.2.9), which was highly found in Seremban and Tampin in Negeri Sembilan. Negeri Sembilan is predominantly a rural state with exceptions in some districts including Seremban and Port Dickson.

The age and sex distribution of the samples also indicated the group of patients most exposed to COVID-19 during the fourth wave of infection. It was clear in our study, that the most affected group age during the fourth wave was 31-40 years, followed by 21-30 years. Notably, at national level, the largest age group contributing to the number of deaths by COVID-19 was 41-59 years, followed by 15-40 years (20). It is possible to deduce that the 31-40 age group is the most vulnerable age to COVID-19 – perhaps due to the working class or increased physical contact largely contributed by this age group (21). Nevertheless, it is important to note some limitations and bias in the sequenced data. For example, elderlies or Orang Asli with limited access to district health centres or hospitals. Therefore, this highlights the thorough and unbiased sampling and sequencing efforts to unravel the evolution and diversity of COVID-19 (22).

Our results show that within the fourth wave of COVID-19 infection, a number of novel mutations circulating within a sub-rural region of Malaysia have emerged. In our dataset, 50 SNPs emerged in 42-417 of the samples and were categorised as high-frequency mutations. These high-frequency mutations were also annotated as classical mutations for the variants Delta and Omicron, respectively. Thus, it was interesting to observe the five novel mutations which also only occurred transiently in our samples. Although SNPs at high frequencies have previously been shown to potentially increase viral transmissibility and virulence (23), detection of short-lived S protein mutations is crucial to detect or predict future mutational sites in emerging variants (24). The comparison of the wild-type protein and mutated model of the spike protein has allowed structural insights of the atomic variation caused by SNPs (25). In our study, none of the five mutations was located in the receptor-binding domain (RBD) and showed no striking structural differences compared to the reference. However, only the V826L substitution occurred in S2 and was more focused in this study due to the high conservation of the S2 region. In the reference model, the 827-826 loops were missing dismissing the direct structural comparison of V826L. At a molecular level, the substitution of L826 could substantially confer increased flexibility to the original loop. Both valine and leucine are aliphatic branched and hydrophobic amino acids. However, valine is slightly more polar than leucine due to the lesser alkyl groups, thus, could confer a more flexible loop (26). Nevertheless, increased flexibility, particularly in loop regions are often compensated by other mutations, thus conserving the protein structure and function. It is also important to note that our study only compared mutations at residual levels, whereas the change of protein structure and function would require whole interprotein interactions.

## Conclusion

Overall, our work has included the spatiotemporal analysis for circulating SARS-CoV-2 lineages at the regional level for Negeri Sembilan. The observed lineages contained high numbers of mutations and variations over a wide range of age groups and districts in Negeri Sembilan – which highlights the rapid evolution of SARS-CoV-2, even at a regional level. While our data simulates what could be the accurate representation of the infection patterns in Negeri Sembilan and, indirectly, Malaysia, our data show that thorough surveillance from all regions of the country is needed for comprehensive management of COVID-19 infections. Nevertheless, from the sequenced samples, we were able to perform molecular and structural analysis to extract novel mutations. Such information can then feed into viral-host interaction experiments to further understand the pathogenesis of SARS-CoV-2.

## Acknowledgements

This study was funded by the Universiti Sains Islam Malaysia Internal Research Grant (PPPI/FPSK/0122/USIM/14322). We also gratefully acknowledge all data contributors, i.e., the Authors and their Originating laboratories responsible for obtaining the specimens, and their Submitting laboratories for generating the genetic sequence and metadata and sharing via the GISAID Initiative, on which this research is based.

## Supporting information

**S1 Table. List of PredictSNP predictions in Negeri Sembilan from July 2021 – May 2022**

## Notes

### Competing Interest Statement

The authors have declared no competing interest.

